# Differences in sodium channel densities in the apical dendrites of pyramidal cells of the electrosensory lateral line lobe

**DOI:** 10.1101/592758

**Authors:** Sree I. Motipally, Kathryne M. Allen, Daniel K. Williamson, Gary Marsat

## Abstract

Heterogeneity of neural properties within a given neural class is ubiquitous in the nervous system and permits different sub-classes of neurons to specialize for specific purposes. This principle has been thoroughly investigated in the hindbrain of the weakly electric fish *A. leptorhynchus* in the primary electrosensory area, the Electrosensory Lateral Line lobe (ELL). The pyramidal cells that receive inputs from tuberous electroreceptors are organized in three maps in distinct segments of the ELL. The properties of these cells vary greatly across maps due to differences in connectivity, receptor expression, and ion channel composition. These cells are a seminal example of bursting neurons and their bursting dynamic relies on the presence of voltage-gated Na^+^ channels in the extensive apical dendrites of the superficial pyramidal cells. Other ion channels can affect burst generation and their expression varies across ELL neurons and segments. For example, SK channels cause hyperpolarizing after-potentials decreasing the likelihood of bursting, yet bursting propensity is similar across segments. We question whether the depolarizing mechanism that generates the bursts presents quantitative differences across segments that could counterbalance other differences having the opposite effect. Although their presence and role are established, the distribution and density of the apical dendrites’ Na^+^ channels have not been quantified and compared across ELL maps. Therefore, we test the hypothesis that Na^+^ channel density varies across segment by quantifying their distribution in the apical dendrites of immunolabeled ELL sections. We found the Na^+^ channels to be two-fold denser in the lateral segment than in the centro-medial segment, the centro-lateral segment being intermediate. Our results imply that this differential expression of voltage-gated Na^+^ channels could counterbalance or interact with other aspects of neuronal physiology that vary across segments (e.g. SK channels). We argue that burst coding of sensory signals, and the way the network regulates bursting, should be influenced by these variations in Na^+^ channel density.

## Introduction

Neurons possess a variety of ion channels and membrane proteins that shape their response properties, from the classical Na^+^ and K^+^ ion channels generating action potentials to G-protein coupled receptors (e.g. Duménieu et al., 2017; Lizbinski et al., 2018; Prešern et al., 2015). Heterogeneity in neuronal physiology can be understood through two complementary principles. One perspective stresses that a given neural output can result from various composition of channels and receptors. This principle was most obviously demonstrated in the stomatogastric ganglion of crabs, where an identical motor output pattern could be generated using networks which neurons differed widely in their channel composition (Prinz et al., 2004). This work highlighted that a change in one element of the neuron’s physiology can be compensated by changes in another element.

The complementary, but non-exclusive, principle is also a basic concept in neuroscience. This principle argues that changes in membrane proteins (e.g. voltage gated ions channels) are necessary for specialization of neurons, and result in different neural outputs (Hille, 2001). An example of this principle, central to the subject of our study, comes from neurons that possesses specific ionic conductances responsible for generating burst firing (Krahe and Gabbiani, 2004). The neuron’s bursting dynamic could not be possible without these specific ion channels and thus their role in neural coding is changed by this bursting dynamic.

Bursting is observed in various sensory systems and typically fulfils the same function: signaling the occurrence of specific spatio-temporal patterns of inputs (Gabbiani et al., 1996; Kepecs et al., 2002). In the visual system, bursts signal edges and sharp contrasts (Lesica and Stanley, 2004), in the cricket auditory system they signal salient ultrasound pulses typical of insectivorous bats (Marsat and Pollack, 2006, 2012) and in the electrosensory system they signal prey-like peaks in signal amplitude or communication signals (Gabbiani et al., 1996; Marsat et al., 2009; Oswald et al., 2004). The presence of a bursting mechanism is key in shaping the neuron’s role in the sensory pathway. The present study focuses on this bursting property and investigates variations in the ion channels responsible for burst generation in the sensory system of weakly electric fish.

The electrosensory lateral line lobe (ELL) is the primary sensory area in the hindbrain of gymnotid weakly electric fish and the main output neurons, pyramidal cells (PCs), possess a well characterized bursting mechanism (Doiron et al., 2002; Turner et al., 1994). PCs, particularly the more superficial ones, have extensive apical dendrites dedicated to receiving feedback inputs, but these dendrites also support the generation of bursts. These apical dendrites extend several hundred μm into the molecular layer (Figure 1A) dorsal to the pyramidal cell layer (Bastian and Courtright, 1991). They contain TTX-sensitive voltage-gated sodium channels (Nav channels). When an action potential is initiated in the cell, the somatic potential backpropagates actively ~200 μm up the apical dendrites due to the Nav conductance. Current from the backpropagating action potential then flows back down to the soma passively causing a depolarizing after-potential (DAP) after each spike (Figure 1B; Turner et al., 1994). In slices and in models, this backpropagation mechanism can trigger a sequence of several spikes with increasingly shorter interspike intervals (ISIs). This stereotyped bursting dynamic, named ghostbursting (Doiron et al., 2002), might not unfold in the same way *in vivo*, where bursts are typically truncated to be only a few spikes-long. Nevertheless, backpropagation and the DAP are an integral part of the bursting dynamic and enable burst-coding of sensory signals (Oswald et al., 2007).

**Figure 1:**
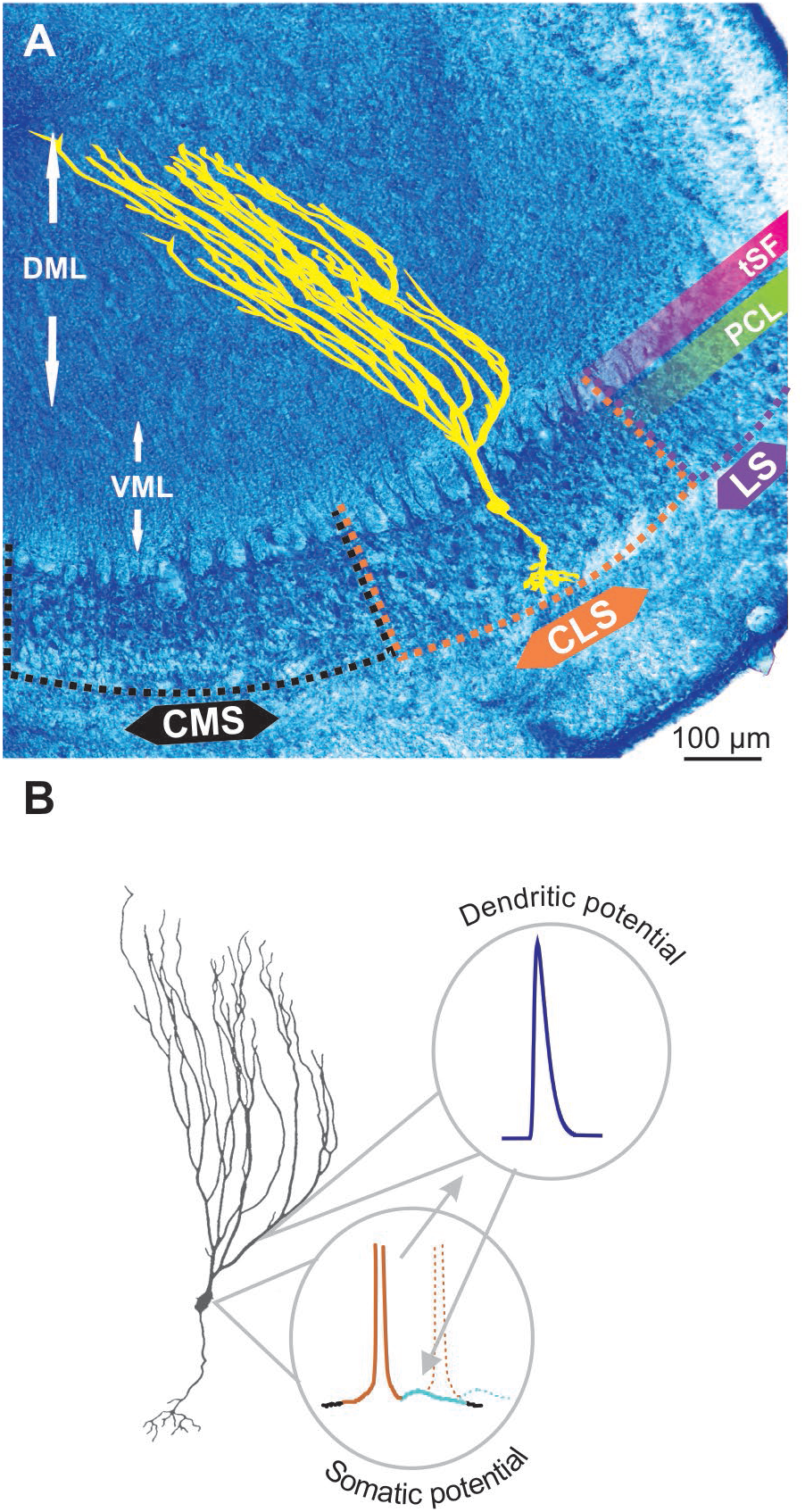
Pyramidal cell and backpropagation. **A.** Transverse section through the electrosensory lateral line lobe (ELL), the primary sensory area located in the hindbrain of gymnotid weakly electric fish. The structure is layered and the main output neurons-pyramidal cells (yellow schematic) are organized in several topographic maps: medial, centro-medial, centro-lateral and lateral segments (MS, CMS, CLS and LS respectively). The latter three segments receive inputs from tuberous electroreceptors. The pyramidal cell layer (PCL) contains the soma of these neurons whereas the ventral and dorsal molecular layers (VML and DML respectively) contain the extensive apical dendrites of the pyramidal cells (illustrated in yellow). The tractus stratum fibrosum (tSF) is an easily identifiable band (see also Fig. 2) separating the PCL from DML. **B.** Backpropagation and bursting. The ON pyramidal cells possess a small basilar dendritic bush receiving feedforward inputs, extensive apical dendrites (particularly long in superficial PCs) receiving feedback and an axon (not depicted here). Apical dendrites contain Nav channels such that, when an action potential is elicited in the soma (brown portion of the voltage schematic), it propagates up and back down the apical dendrites causing a depolarizing after potential (DAP; light blue) at the soma. The DAP increases the probability that subsequent action potentials (e.g. dashed line) are fired immediately after the first one thus leading to bursting.

Response properties of ELL PCs are shaped by a variety of additional factors that vary across cell subtypes (Chacron et al., 2011; Maler, 2007). PCs are classified based on their location in the ELL layer. Superficial and intermediate PCs have extensive apical dendrites while deep PCs have short apical dendrites and fulfill a different role in the circuit (Bastian et al., 2004; Bastian and Nguyenkim, 2001). PCs receive inputs from receptors either directly or through an inverting interneuron. This leads to response patterns typical of ON-cells and OFF-cells respectively (Clarke et al., 2015; Maler, 1979). Both types have the same burst dynamic. The ELL is organized in several topographic maps. While the map of the medial segment is driven by a separate category of passive ampullary electroreceptors, the centro-medial, centro-lateral and lateral segments (CMS, CLS and LS respectively) receive inputs from the tuberous receptors sensitive to the fish’s self-generated electric signal (Kawasaki, 2005). These areas each play a critical role in prey detection, communication, and navigation. In this study, we focus on those three ELL maps responsible for processing this actively-generated electrosensory signal.

PCs from the three maps vary widely in their response properties. These differences are due to variations in network connectivity, ion channels composition, expression of neuromodulator receptors and more (Ellis et al., 2008; Krahe and Maler, 2014; Maler, 2009). These differences in properties allow the specialization of the three maps for different purposes: whereas the CMS is well suited for the localization of small near-by objects such as prey, the more lateral segments (CLS and especially LS) might specialize for processing communication signals (Marsat et al., 2009; Metzner and Juranek, 1997). Despite these differences in properties and function, all segments display similar bursting rates (Krahe et al., 2008). Although some small differences in burst patterns have been noted (Mehaffey et al., 2008; Metzner et al., 1998; Turner et al., 1996), it is unclear to what extent burst dynamics varies across segments since no fundamental differences have been documented in burst coding (Krahe et al., 2008; Metzner et al., 1998; Oswald et al., 2004). Large differences, particularly in bursting rates, could have been expected across different segments given the known differences in conductances affecting bursting. SK channels, which generate a hyperpolarizing after-potential, vary in expression across PCs subtypes and segments. They are particularly prevalent in the LS where they oppose the DAP-based burst generation mechanism (Ellis et al., 2007a, 2008). Serotonin receptors are also expressed differently, with LS cells expressing more, and when activated serotonergic inputs can enhance bursting propensity (Deemyad et al., 2011, 2013; Johnston et al., 1990). It is clear that many factors interact to shape bursting and it is possible that similarities in burst coding across PC subtypes and segments happens despite differences in intrinsic configurations, rather than because they have identical physiology.

Variation in one element central to the burst-generation mechanism has not been examined yet: Nav channel expression in the apical dendrites of PCs. Considering the differences related to the burst mechanism noted above, we hypothesize that variations in the expression of Nav channels will also be observed across segments. To test this hypothesis, we performed immunocytochemistry on ELL slice labelling Nav channels in the molecular layer of the three segments of the ELL. We observed Nav expression throughout the molecular layer and we show that it is denser in LS than CMS. We argue that this differential expression should have functional consequences on response properties but that it is hard to determine how it interacts with the many other differences seen across segments.

## Materials and methods

### Brain slices preparation

*Apteronotus leptorhynchus* fish used for experiments were wild-caught and purchased from a tropical fish supplier. They were maintained in home tanks (61×30.5×50.8 cm) at 26–27°C, 250–300 μS on inverted light cycles, fed *ad libitum* and were provided with environmental enrichment. Fish of either sex were anesthetized in tank water with MS-222 (3-amino benzoic acid ethyl ester, Western Chemicals Inc.) and respirated with oxygen bubbled MS-222 water during perfusion. All chemicals were obtained from Fisher scientific (Hampton, NH) unless otherwise noted. Heart was surgically exposed and intracardial perfusion was performed via the Conus arteriosus with 5 ml of cold 0.9% saline containing Heparin (# 9041-08-1), NaNO2 (# S25560) and NaCl (# 7647-14-5) which is followed by perfusion with 40 mL of cold 4% paraformaldehyde (Electron Microscopy Sciences, #RT-15714) in 1X-phosphate buffered saline (PBS), pH-7.3. Whole brains were surgically removed and post fixed in 4% paraformaldehyde (PFA) in 1X PBS for 4 hours at 4°C and were washed three times for 15 minutes each in 1X PBS at 4°C. Brains were sequentially cryoprotected in 20% and 30% sucrose (# S25590) in 1X-PBS, pH-7.3 until they were completely saturated and later incubated in 1:1 mixture of 30% sucrose solution and optical cutting temperature (OCT) compound (Electron Microscopy Sciences, #62550-01) for 1-2 hours before embedding in OCT. Dry-ice chilled 100% ethanol was used to freeze the brain in OCT in a cryomold and the mold was incubated at −80°C for 1-2 hours before sectioning. 15-20μm thick true-transverse brain sections were obtained using cryostat (Leica 1850) and the slides were stored at 4 °C for immediate processing or stored at −20°C until use.

### Immunohistochemistry

Brain sections were immunoreacted for Nav *in situ* through the following procedure. Sections were washed 3 times with 1X-PBS, pH-7.3 for 5 minutes each and were blocked for 1 hour in 5% normal goat serum (# 005-000-121, Jackson Immuno Research) in PBSAT (1X PBS, 5mM Sodium Azide and 0.1% Triton X-100). Blocking was followed by 1-hour incubation with Anti-Pan Nav (Alomone labs, # ASC003) (1:50) and purified Mouse Anti-MAP II (BD biosciences, #556320) (1:400) primary antibodies in blocking buffer at room temperature. The Anti-Pan Nav antibody was raised in rabbits, and was shown to selectively bind to the Na channels in apteronitids’ electric organ with a protein size of 250-260 kDa (Ban et al., 2015). Later, brain sections were transferred to 4 °C for overnight incubation. Note that the MAP2 antibody used in the current study only stains the high molecular weight isoforms of MAP2 and does not recognize low molecular weight MAP2 isoforms or other microtubule proteins. In addition, MAP2 is mainly concentrated in the dendritic part of the nerve cells (Olesen, 1994), this might possibly explain the comparatively fainter MAP2 labelling observed in the cell bodies.

Sections were washed 4 times with 1X-PBST (1X PBS and 0.1% Triton X-100), pH-7.3 for 15 minutes each and were incubated with Goat anti rabbit Alexa 488 (Life Technologies, #A-11008) (1:500) and Goat anti Mouse Alexa 546 (Thermofisher, #A-11030) (1:500) secondary antibodies for 3 hours at room temperature in an enclosed moist chamber. Sections were washed 4 times with 1X-PBS, pH-7.3 for 15 minutes each and were mounted in Vectashield (Vector Laboratories, # H-1000) and coverslipped. Selectivity of the labelling was confirmed with several controls: an absorption control where the Nav antibody is incubated with the immunogen (Supplementary figure 1), a control with no primary antibody (Supplementary figure 2) and a quantification of Nav labelling in regions where no Nav channels are expected to be found. For this last control, we selected 27 different 10×10um areas in the gaps between dendrite in the tSF layer (where we do not expect Nav channels to be present) and compared the Nav labeling to 27 areas selected randomly on the same scans but in the middle of the dendritic arbors in the molecular layer. As expected, minimal staining was found (gaps in tSF: 0.009 puncta/μm2; molecular layer: 0.1 puncta/μm2; T-Test: p<10^−6^) proving that puncta of labeling are not simply an artefact and randomly distributed on the tissue.

### Western blot

The specificity of polyclonal Anti Pan-Nav antibody was evaluated using western blot analysis (Figure 3F). Unperfused whole brain was surgically removed and freezed in liquid nitrogen and immediately homogenized with cold homogenization buffer containing 250mM Sucrose, 1mM EDTA, 10mM Tris HCL and protease inhibitor cocktail (#4693132001, Sigma Aldrich), pH-7.2, on pre-chilled mortar and pestle. Tissue sample was sonicated using two 10 second pulses with 30 second interval between each sonication. Sample was centrifuged at 500X g for 10 minutes to remove intact cells, nuclei and cell debris, and the supernatant was centrifuged at 100000 X g for 1 hr at 4 °C. Supernatant was discarded, and the pellet was resuspended in homogenization buffer and centrifuged at 100000 X at 4 °C for 1 hour. Resultant supernatant was discarded and the pellet containing the membrane fractions was used to run western blot.

For western blot, 1X LDS (#NP 0008, Thermofisher) was added to the protein samples which was then heated for 10 min at 70°C. 5uL of the sample was loaded into 4-6% polyacrylamide gel along with Precision Plus Protein™ Dual Color Standard (#1610374S, Bio-Rad) and was run at 120 V. Transfer was done at 100mA for 22 hrs at 4 °C and the nitrocellulose membrane (# 162-0094, Bio-Rad) was blocked for 1 hour with 5% BSA and 0.05% NaN 3 in 1X TBST (1X TBS, 0.1% Tween) under agitation. Primary antibody diluted in the blocking buffer (1:200) was applied to the membrane and incubated overnight at 4 °C under gentle agitation. After incubation, membrane was washed 3 times with 1X TBST for 15 minutes each. HRP conjugated anti-rabbit secondary antibody (#A0545, Sigma) was applied at 1:10,000 dilution and agitated for 1 hour at room temperature, which was followed by three 15-minute washes with 1X TBST. ECL substrate (#RPN2109, GE Healthcare) was added and the membrane was imaged on FluorChemQ system (Protein Simple).

### Nav puncta density quantification

Scans were obtained on FV-1000 Fluoview confocal microscope and minor brightness adjustments were made using Fluoview software. All of the scans were imaged using 60X oil immersion lens at 2X digital zoom unless otherwise noted. Scan size X*Y is set at 105.4*105.4μm with Z at 0.5μm. Differences in the spatial distribution across the brain maps is assessed by quantitative image analysis using VAA-3D software. Image J software was used to perform image normalization.

To measure the spatial distribution of dendritic Nav channels across different ELL maps (LS, CLS and CMS), the dorsal edge of the tractus stratum fibrosum (tSF) layer is chosen as reference location (0 μm). Scans covered the first 600 to 800 μm of apical dendrites as they project dorsally through the VML and DML layers.

In each scan, 2-4 portions of dendrites with clear MAP2 labeling were chosen prior to looking at the Nav labelling to prevent biased choice of dendrite based on expected channel density. Nav labeling was quantified by manually marking each punctum visually identified on the chosen dendrites. XY coordinates of each punctum and dendrites were stored for analysis. We were conservative in assigning a punctum as belonging to a dendrite and the numbers presented in this paper should be viewed as a lower-bound estimate. Note that any bias in quantifying puncta within a segment is also inherent to the quantifications done across segments, and therefore, the differences observed in Nav density across the segments are unlikely to be affected by such biases.

To standardize the identification of the edge of the tSF layer (defined as location 0 μm) a Matlab (Mathworks, Natick, MA) program was created to plot the density profile of the gaps present in the tSF layer along with the pixel intensity profile of the scan (see Figure 2D and explanation in the legend).

**Figure 2:**
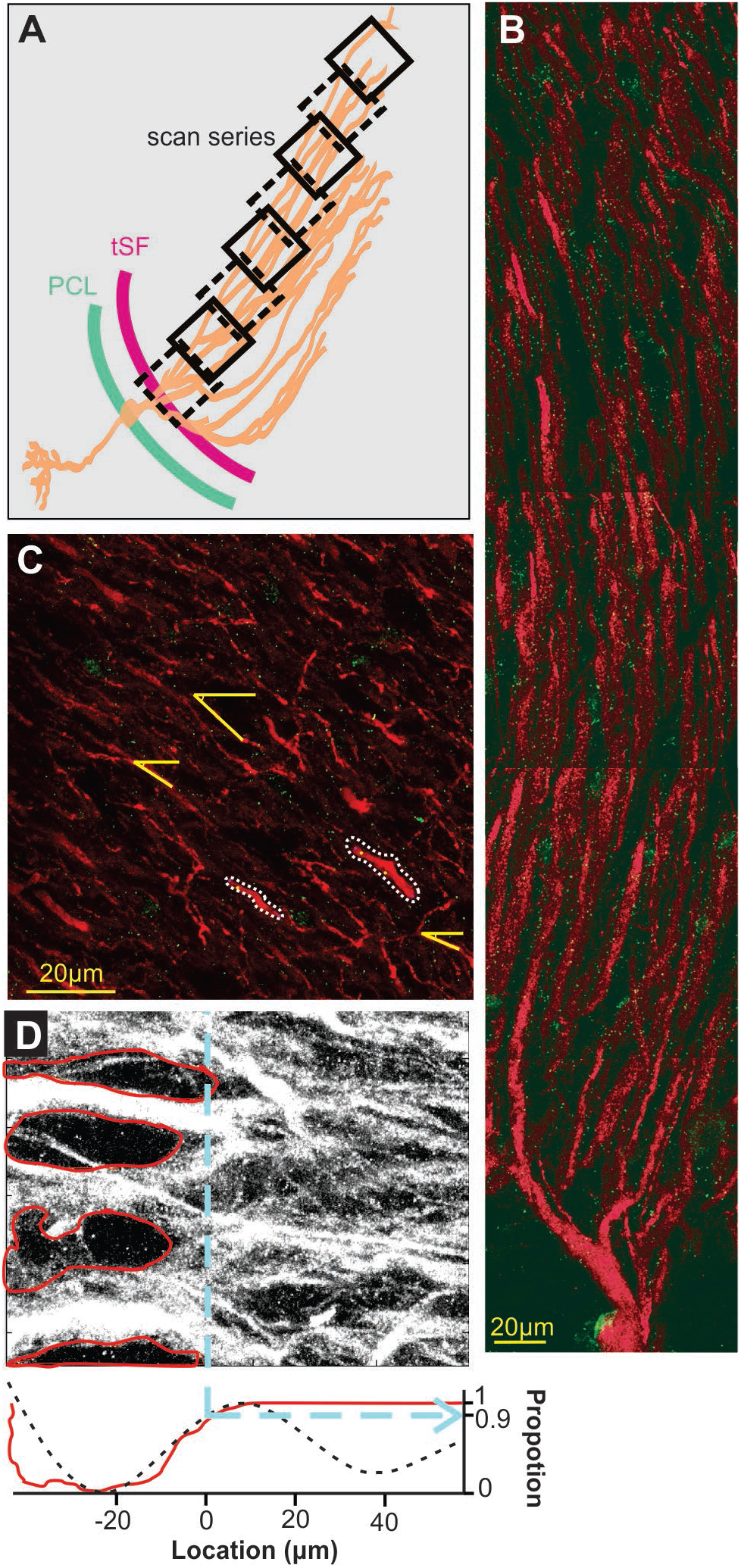
Scanning and localization procedure. **A.** Scans arrangement. Each 105×105 μm scan is positioned relative to the transverse section of the ELL so that the first one of a series contains the tSF layer and subsequent ones overlap the previous and follow the orientation of dendrites in the molecular layer. Eight successive scans or more were performed to cover at least 600 μm of apical dendrites. **B.** A series of scans showing the extensive apical dendritic bush in the molecular layer. Although we did not attempt to reconstruct and isolate single pyramidal cell, our successive scans follow the apical dendrites of PCs. Red MAP2 labelling marks the inside of the dendrites and green labelling highlights the position of Nav channels. Since no distinction could be made between ON and OFF cells based on the immunolabelling used, all the dendrites quantified in the study are mixed, either ON or OFF-cells, and possibly VML interneurons (although it is unknown if they express dendritic Nav). **C.** Dendrite selection and angle. Based solely on the MAP2 labelling (Nav labeling not displayed) we selected several clearly defined dendrites per scans for quantification (e.g. outlined with white dashes). Since dendrite orientation relative to the scan orientation is not orthogonal, we measured, in each scan, the average relative orientation of dendrites based on several dendrites per scan (yellow lines). This allowed us, using the position on the x-axis of the scan, the angle of dendrites and trigonometry, to determine a more accurate position of dendrites along the dendritic bush. **D.** Determination of tSF dorsal edge to set it as location “0μm”. The stratum fibrosum tract is characterized by large circular areas without MAP2 labelling between the large proximal apical dendrite shafts. We determined the dorsal edge of this layer by first highlighting the visible holes in labelling (red in the top image) and constructing a pixel histogram (red curve, bottom plot) along the x-axis based on the pixels outside these areas. Second, we used the raw pixel intensity values to build a histogram along the x-axis and smoothed it by fitting a triple sinusoid function (black dashes, bottom plot). Both histograms were normalized between 0 and 1 and we took the 0.9 mark of the rising slope to determine the 0 μm location (dashed blue line). We confirmed both histograms gave similar results and used the average of the two calculated values to set the 0-mark.

A series of slightly overlapping scans of the entire extent of each map were obtained (Figure 2A) and the bleached area from the overlapped portion as well as a local landmark was used to obtain the start point x-coordinate of each subsequent scan of the map. This allowed us to determine the orthogonal distance of each scan, dendrite and punctum from the tSF edge. Since dendrites do not travel orthogonally to the tSF edge and the scan orientation (Figure 2B&C), we had to correct the distances using dendrites’ angles. To do so, in each scan we selected randomly 5 clear portions of dendrites and measured their angle relative to the scan’s frame (Figure 2C). Using the average angle of the selected dendrites in each scan and trigonometry, orthogonal distances could be converted into estimated distance (i.e. distances that take into account the general curvature of dendrites). This estimate is still an underestimation of actual distance along the dendritic tree since dendrites are often not completely straight over one scan, and also any z-plane curvature was ignored. Note that we sectioned the hindbrain in a true-transverse orientation minimizing the curvature of dendrites in the z-plane over the proximal apical dendrites. This underestimation does not affect our conclusions but should be noted. Data analysis was performed with Matlab (Mathworks, Naticks, MA) and statistics with JMP (SAS Institute Inc, Cary, NC).

### Burst ISI characterization

Fish were prepared for in vivo electrophysiological recordings as described in Marsat et al., (2009) where recording and analysis methods are also described in detail. Briefly, superficial ON and OFF-cells of the three segments were targeted and spontaneous activity was recorded for 60 seconds. Recorded spike trains were binarized and burst identified by first constructing an ISI histogram of the entire spike train and identifying the upper interval limit characteristic of the cell’s burst ISIs (see Marsat and Pollack, 2012 for more details). Identified bursts were then used to calculate their ISI distribution. All procedures on animals were authorized by West Virginia University’s IACUC (protocol #13-0103).

## Results

To study the distribution of Nav channels along the apical dendrites of PCs we used a pan-Nav antibody known to bind selectively to these channels in other tissues of this species (Ban et al., 2015). We confirmed the selectivity of the antibody in brain tissue by performing a Western blot (Figure 3F). To identify the position of the labeled channels relative to the dendrites, we also used a MAP2 antibody that densely labels the microtubules inside dendrites (Deng et al., 2005). By using thin brain slices (15-20 μm) and high-resolution confocal scans we were able to precisely localize the labeled channels relative to PCs apical dendrites (Figure 3A-E). The pattern of expression of Nav channels in the dendrites followed a punctate pattern, as described previously (Turner et al., 1994), that is visibly different from the more uniform expression pattern in the soma of interneurons (Figure 3B) or in axons. Prior to visualizing the Nav labelling, we selected in each scan 2-4 portions of dendrites that are clearly delineated by the MAP2 labelling. We then visualized in 3D (moving through the z-plane with the imaging software) the position of each punctum in the vicinity of the dendrite portion selected for quantification. Since Nav channels are in the membrane of the dendrites and MAP2 proteins are inside the dendrite, Nav puncta were immediately adjacent (in the x-y plane or the z-plane) to the MAP2 labelling or overlapping.

**Figure 3:**
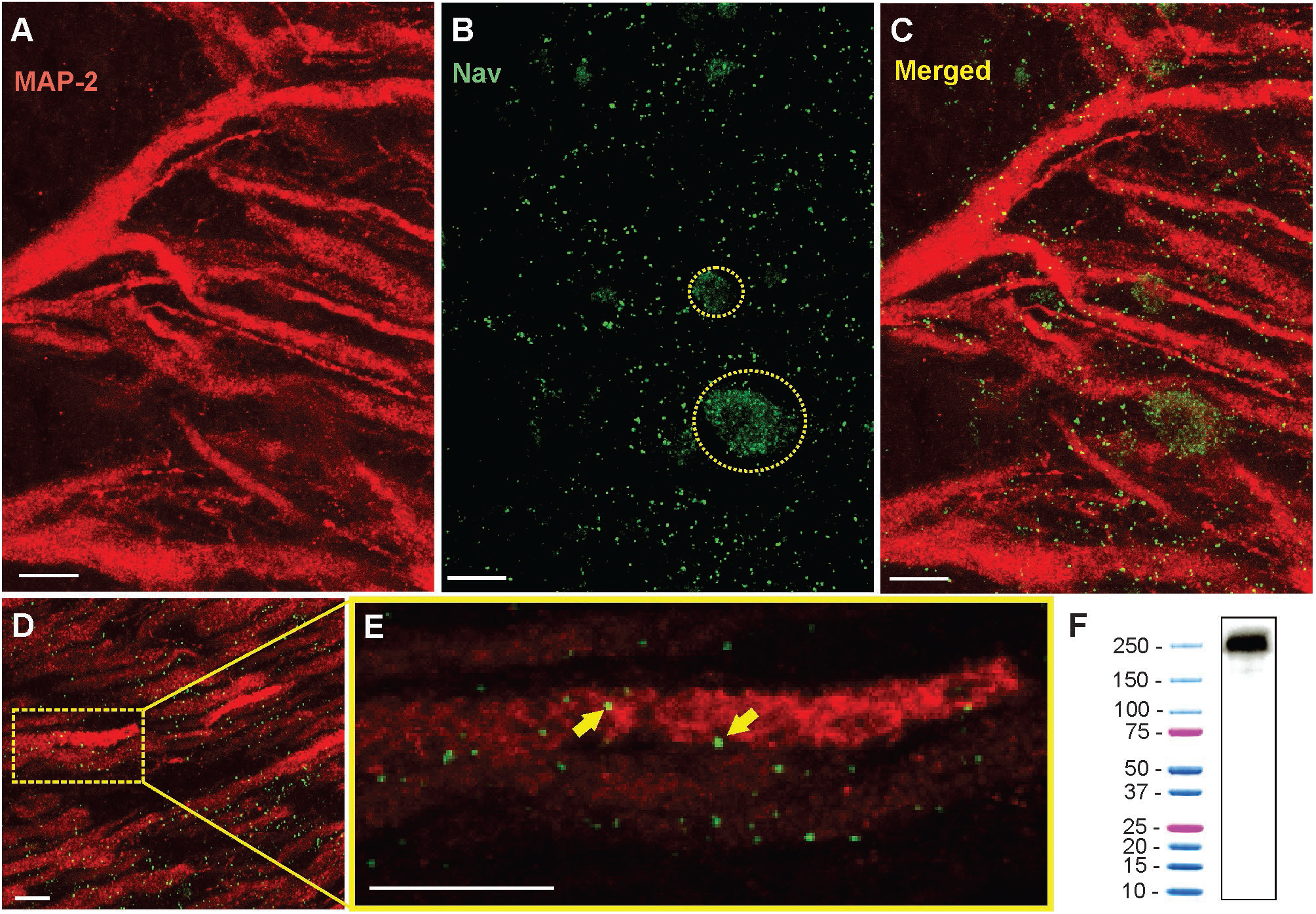
Immunohistochemistry show the punctate expression of Nav channels on the apical dendrites. **A.** Confocal image of a 16 μm section of ELL showing the red Anti-MAP2 labeled apical dendrites projecting through tSF and VML. White scale bars are 10 μm in all panels. **B.** Nav channel punctate distribution (green dots). Nav label was also found on the surface membranes of interneurons (e.g. VML cells) where their distribution is more uniform. Two interneuron cell bodies are outlined with a dashed yellow line. **C.** Merged MAP2 and Nav labelling. Note that these images show the combined z-layers of the scan through the 16 μm slice but that looking through the individual successive scans at varying depth (0.5 μm apart) can help determine the proximity between MAP2 and Nav labelling. **D.** Nav and MAP2 labeling in a more distal portion of VML (~200 μm from tSF boundary). **E.** Enlarged view of one dendrite in the image displayed in D selected for quantification of Nav channels. The proximity between the labelled Nav channels and the labeled MAP2 inside dendrites allows us to confirm that the channels were in the membrane of a pyramidal cell dendrite. Two puncta are indicated with yellow arrows. Note that the Nav and MAP2 labelling does not need to overlap directly since MAP2 is located inside the dendrite and Nav channels are in the membrane. We thus expect some of the puncta to be slightly separated (in the x-y plane or the z plane) from the MAP2 labelling. **F.** Western blot analysis of brain tissue demonstrating the specificity of Anti-pan Nav antibody. The western blot of the tissue processed along with a protein ladder displayed a single band at ~250-260 kDa.

Our dataset is based on images from 15 brain slices (5 slices each from 3 fish). In each slice, all three segments were scanned and quantified thereby assuring that differences in immunolabeling clarity from slice to slice could not cause a bias in quantification across segments. Scans starting at the tSF layer and tiled to extend beyond 600 μm into the molecular layer (Figure 4) allowed us to evaluate Nav expression as a function of segment and location along the ventral-dorsal axis (i.e. proximal-distal to the soma). We localized and counted Nav puncta on 727 portions of dendrites representing over 21400 μm of dendrites (Figure 5). We found a large dendrite-to-dendrite variability in puncta density with some long portions of dendrites containing no puncta and others being densely populated with puncta. This variability could be observed even between dendrites located side-by-side on the same scan precluding the possibility that variations in the immunolabeling process could account for this variability.

**Figure 4:**
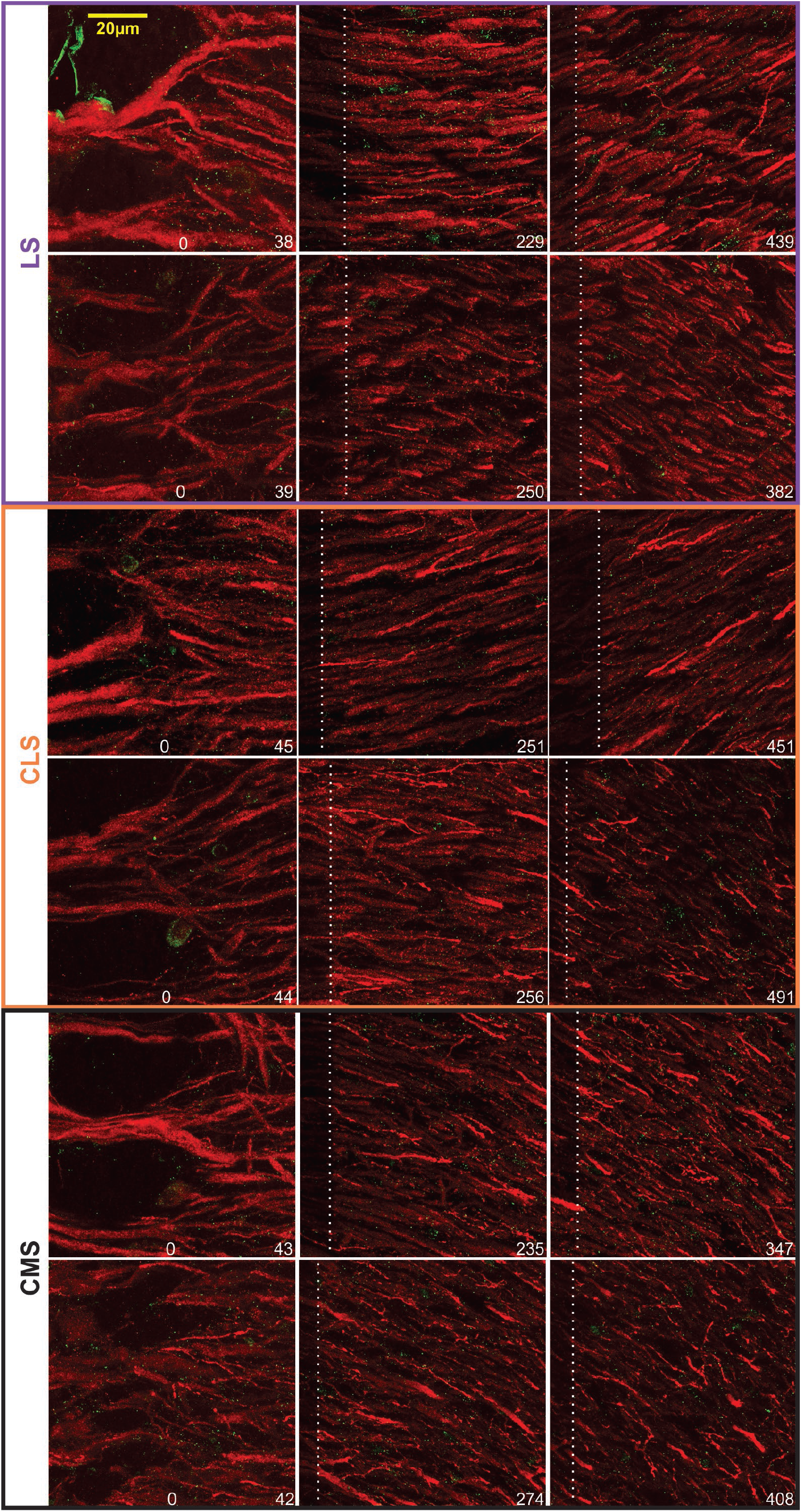
Examples of Nav-Map 2 expression in the ELL. Confocal images from 2 brains (top vs bottom rows for each map) taken at different distances from the tSF. The location of the distal edge of the image is specified in μm at the bottom right of each image. The white dashed lines indicate the edge of the previous -overlapping-scan and thus the pixels on the left of this line are bleached from the previous scan.

**Figure 5:**
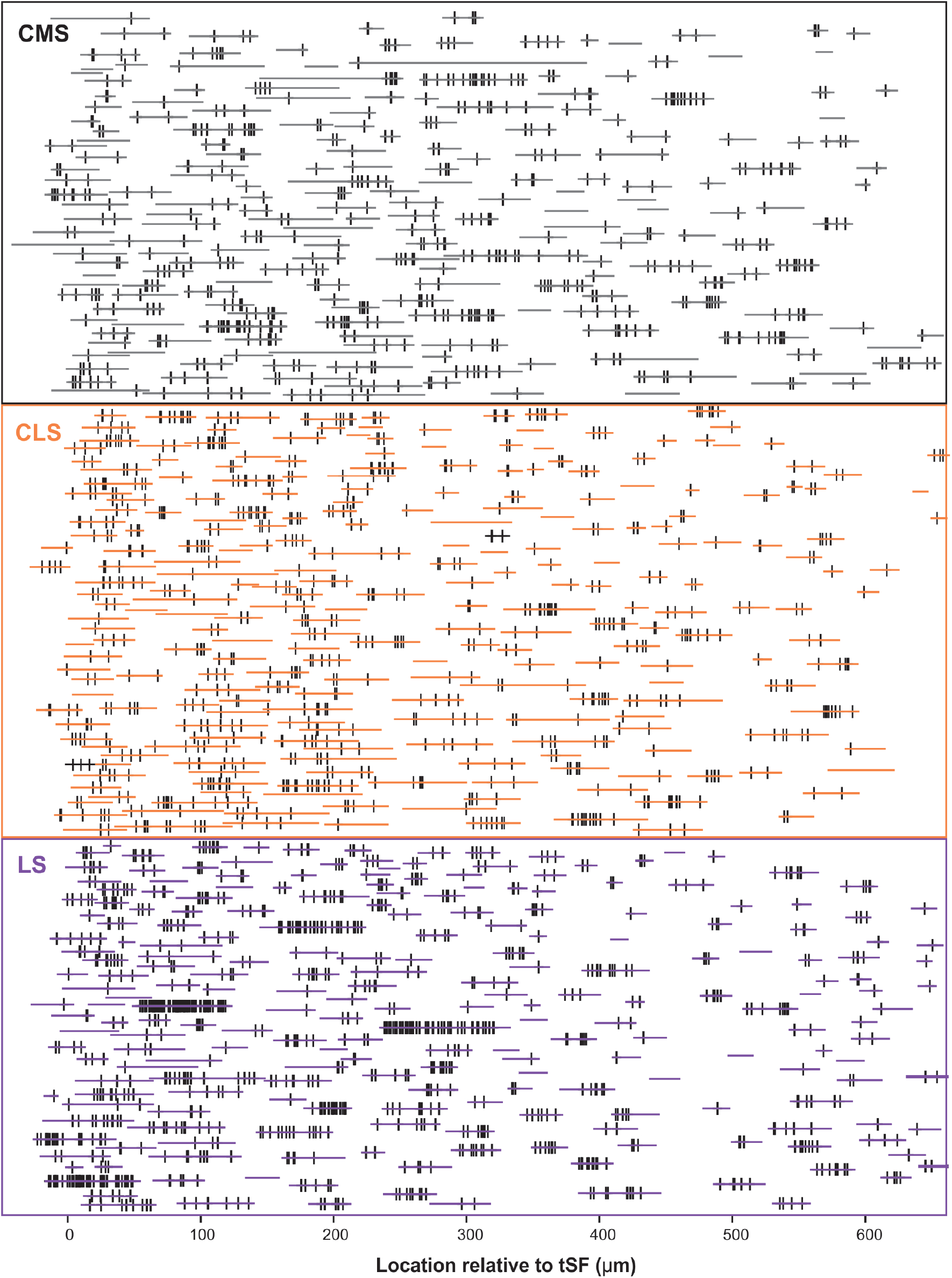
Schematic representation of the entire dataset. The position and length of each dendrite quantified is depicted by the horizontal colored lines. The position of each Nav punctum identified are marked by a vertical black line. The vertically stacked arrangement of the dendrites loosely follows the order of processing and thus data from a given slice/brain are found in adjacent rows of the stack. We quantified 727 dendrites in 15 brain slices, totaling >21400 μm of dendrite, and identified 2435 puncta of Nav labelling across segments/locations.

PCs can be classified in several subcategories (deep, intermediate or superficial PCs and ON-cells or OFF-cells) organized in microcolumns and their apical dendrites can overlap at the same location in the molecular layer. Our method cannot determine to which type of PC a dendrite belongs and thus we cannot determine to what extent the variability observed is due to variations across -vs. within-subcategories.

The density of puncta per μm of dendrite was calculated for each of the ~250 dendrites per segment and we first visualize in a 3D histogram the proportion of dendrites with various densities as a function of dendrite position along the molecular layer (Figure 6A). The pattern of Nav expression clearly differs across segments. In LS, a large proportion of dendrites have puncta densities higher than 0.1 puncta/μm whereas CMS dendrites have densities mostly below 0.1 puncta/μm. Our data also shows that density stays similar throughout the molecular layer up to at least 600 μm away for tSF. Our data beyond 600 μm are more sparse, nevertheless, the data we do have suggest that more distal dendrites express Nav channels with a density similar to the more proximal dendrites (see Figure 5 and 6A). Our data demonstrates that dendrites in the LS have Nav puncta densities about twice as high as CMS dendrites and 1.4 times higher than CLS dendrites (Figure 6B-C). The distribution of Nav puncta along the first 600 μm of dendrites (for which we have most data) was not very different across segments. Specifically, we see similar densities over the length of the dendrites in LS and CLS but channel density increased slightly in CMS for dendrites located further from tSF (Figure 6D).

**Figure 6:**
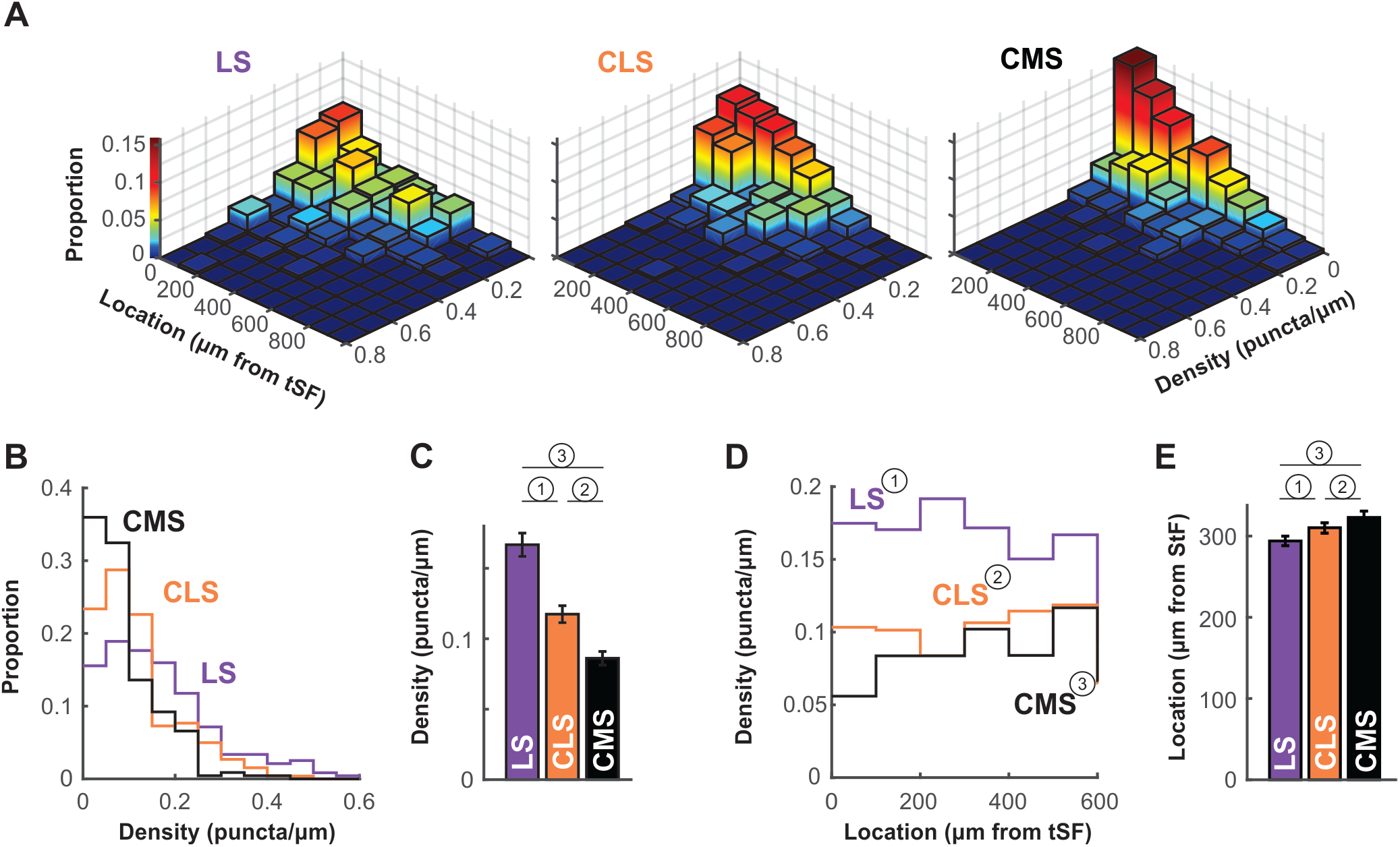
Nav channel expression density differs across maps. **A.** Histogram of channel density per dendrite as a function of location. For each dendrite (LS, n=238; CLS, n=261; CMS, n=228) channel density (puncta/μm) was calculated. The z-axis (also color-mapped) shows the proportion of dendrites in each location/density bin (i.e. sum of all bin heights is 1 in each panel). These data show that densities in CMS are largely constrained to the 0-0.1 puncta/μm range whereas densities are equally distributed in the 0-0.1 and 0.1-0.2 range (and higher) for LS (see panels B-C). Also, channel density seem to be equally distributed along at least the first 600 μm of the molecular layer (see panels D-E). Note that overall height of the bins tends to decrease with increasing distance from tSF simply because fewer dendrites were quantified distally compared to proximal locations. **B.** Histogram of channel density across maps. The data displayed in A were collapsed across all locations to clearly show the differences in puncta density across maps. LS clearly has a higher proportion of dendrites with high density (>0.1 puncta/μm) compared with CMS and thus a lower proportion of dendrites with low density (<0.1 puncta/μm). See panel C for a listing of statistical differences. **C.** Average puncta density is significantly different across segments and decreases from LS to CMS. Since the data are not normally distributed (see panel B) we tested pairwise differences using a Wilcoxon test: (1) p=0.000004; (2) p=0.0002; (3) p=10^−15^. **D.** Channel density is similar across location in the molecular layer. Average channel density was calculated for dendrite positioned at various distances for tSF. Location does not influence the observed puncta density in LS and CLS (One-way ANOVA: (1) p=0.82; (2) p=0.1) but density increases slightly with distance in CMS (One-way ANOVA: (3) p=0.005). The analysis shown in this panel and in panel E took in account dendrites located <600 μm only since our data for locations >600 μm is sparse. **E.** Nav channel distribution across the molecular layer is similar for all three maps. We calculated an average channel location across the 600 μm of the molecular layer considered here to compare the distribution of channels along the dendrites for the three maps. To do so, we normalized our puncta counts to account for the length of dendrites quantified in each location bin. Therefore, an average location of 300 μm indicates that channels are uniformly distributed across the dendritic length considered here. This average is similar for all three maps ranging from 294 μm to 323 μm but the small differences are significant (Wilcoxon: (1) p=0.0001; (2) p=0.003; (3) p=10^−11^).

Therefore, our data shows that, despite small differences across segments, we see a qualitatively similar distribution of Nav puncta over the entire length of dendrites considered here (0-600 μm from tSF dorsal edge; Figure 6E).

The differences in Nav channel density across segments could have a significant impact on the neuron response properties. The impact on neural dynamic and sensory processing is hard to gauge because PCs differ in many aspects between segments. Differences in ion channel composition, tuning, connectivity and more, interact intricately with the dynamics imposed by Nav channels. Even potential studies using modeling, where a single parameter can be altered (e.g. Nav channels density), presents important challenges. Indeed, all the elements of the neurons and of the network known to influence the neuron’s response dynamic would need to be included to understand the effect of changing Nav density and determine if it can explain differences observed across segments. Such modelling effort is beyond the scope of this manuscript.

Nevertheless, we suggest that looking at spontaneous activity of the neurons could give us useful insight into the effects of differences in Nav density on bursting. Interspike-intervals (ISIs) during spontaneous activity are largely determined by the neuron’s intrinsic mechanisms since excitation is weak and relatively constant. Furthermore, burst ISIs are heavily influenced by only a few ion channels, first and foremost the Nav channels in apical dendrites. Therefore, we investigated potential differences across segments in burst ISIs during spontaneous activity. Focusing on superficial pyramidal cells, we show that burst ISIs tend to be shorter in LS than in CMS; CLS being intermediate (Figure 7; average ISIs ± s.e.: LS= 3.2± 0.6; CLS= 3.8± 0.5; CMS=6.2± 0.3. Wilcoxon test: LS-CLS, p<0.0001; LS-CMS, p<10^−9^; CLS-CMS, p<10^−7^. n=13 to 19 neurons). This trend is consistent with a previous observations (Mehaffey et al., 2008; Turner et al., 1996) and could potentially be attributed in part to the higher density of Nav channels in LS (see discussion).

**Figure 7:**
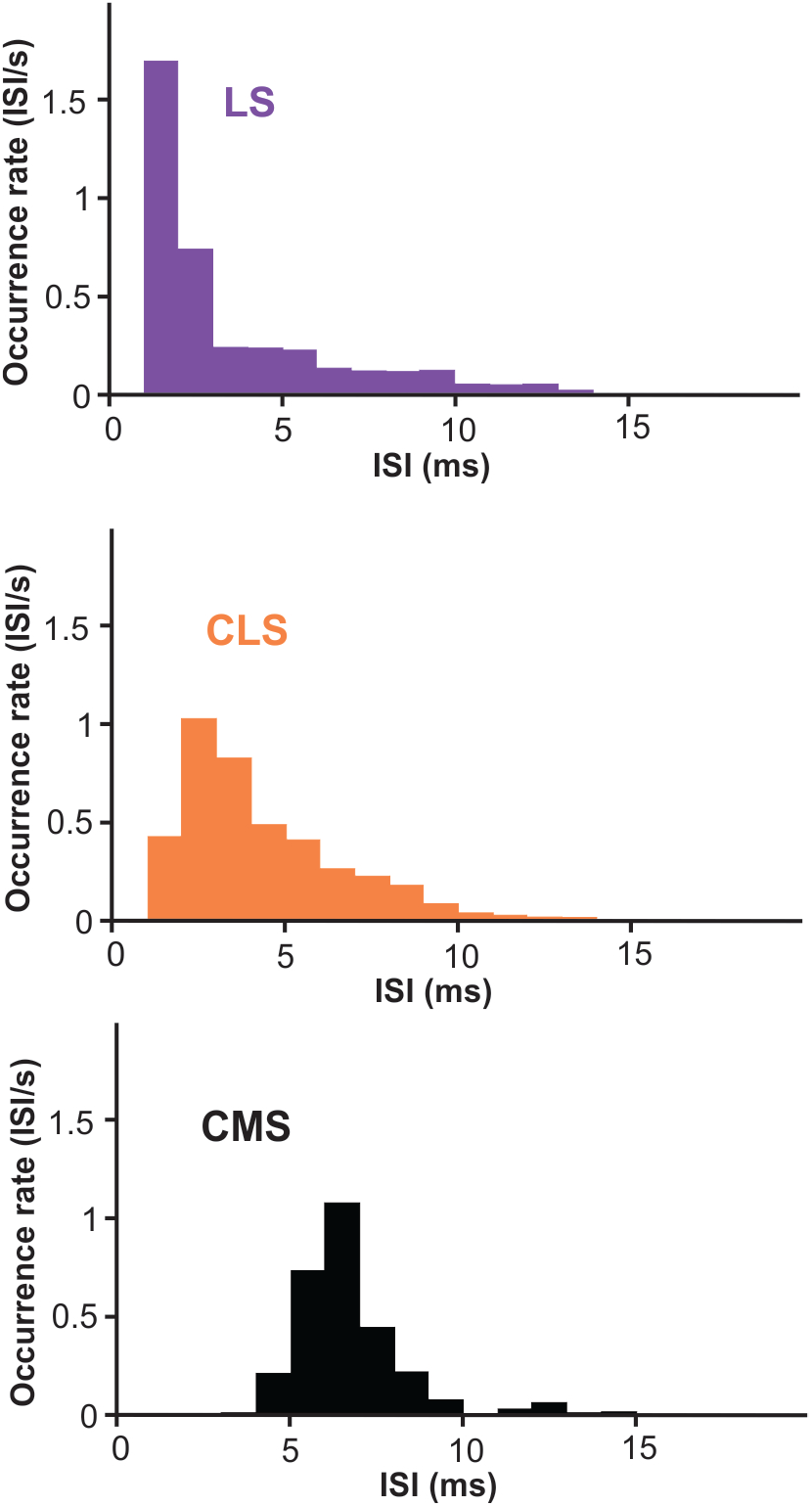
Inter-spike intervals (ISI) histogram for burst-spikes during spontaneous activity. Burst of spikes were identified as described in the methods and the interval between the spikes in each burst contributed to the histograms displayed. The average interval is shorter in LS than CMS with significant differences across all three segments (averages ± s.e.: LS= 3.2± 0.6; CLS= 3.8± 0.5; CMS=6.2± 0.3. Wilcoxon test: LS-CLS, p<0.0001; LS-CMS, p<10^−9^; CLS-CMS, p<10^−7^. n=13 to 19 neurons).

## Discussion

### Dendritic Nav channel expression and firing patterns

The principle finding of the present experiment was that LS exhibits the highest dendritic Nav channel density, followed by CLS and CMS. The difference was obvious in the proximal dendritic regions of apical dendrites that support active backpropagation. Nav channel densities remain relatively constant along the proximo-distal axes of the apical dendrites in all segments.

Previous studies have established the subcellular distribution and function of TTX sensitive Nav channels using immunocytochemistry and electrophysiology techniques (Turner et al., 1994). Punctate regions of Nav immunolabel were detected in pyramidal cell somata, basal and apical dendrites. The previous EM study (Turner et al., 1994) pointed out that regions of denser labelling along a dendrite could be separated by longer portions of dendrites with little labelling. Our study confirms that channel density varied widely from one portion of dendrite to the next. Note that our study does not distinguish between dendrites belonging to ON-cell, OFF-cells or interneurons. Part of the variability observed could be due to differences across cell types If this was the case, you would expect to see 2 or 3 clusters of channels densities (one for each cell types) but we observe a mono-modal distribution instead. Future studies labelling for cell identity and for Nav could test this hypothesis.

Dendritic Nav channels allow the active propagation of an antidromic spike over around 200 μm of proximal apical dendrite. The resulting DAP at the soma after each somatic spike underlies the production of burst discharge. Focal ejections of TTX inactivates Nav channels, decreasing spike frequency (Oswald et al., 2004; Turner et al., 1994). However, higher conductance from Nav channels does not necessarily lead to an increase in spike frequency. Decreased Na^+^ conductance in dendrites can increase excitability of the soma by delaying the DAP thereby enhancing the so-called “ping-pong” dendro-somatic dynamics that underlies burst generation (Fernandez et al., 2005). There is thus a non-monotonic relationship between Na^+^ conductance and cell excitability (Fernandez et al., 2005) or the amount of information carried by bursts (Doiron et al., 2007) with a maximum at intermediate values. Nevertheless, modeling studies indicate that an increase in dendritic conductance from Nav channels systematically causes increased burst rates (Doiron et al., 2007).

Backpropagation has been characterized as traveling up 200 μm before the active propagation is not detected. Our results, and the original study that identified the Nav channel’s presence in the dendrites (Turner et al., 1994), show that the channels are distributed along the majority of the dendritic tree. The role of these channels, past the 200 μm where backpropagation is evident, is unknown. One possibility is that they contribute to the DAP but the current they generate in the more distal dendrites is too small to be clearly identified. The DAP is indeed likely to be reduced in distal apical dendrites due to the presence of Kv3.3 channels (Deng et al., 2005) producing an after-hyperpolarization. Another possibility involves an interaction between the channels and synaptic inputs. GABAergic inputs have been shown to influence the DAP generation (Mehaffey et al., 2005), it is therefore not unlikely that the Na^+^ current in the distal dendrites shapes and affects the dynamics of synaptic inputs. Given the important role of feedback inputs onto these apical dendrites the presence of the channels near these synapses could alter the neuron’s response properties significantly.

### Segment-specific regulation of bursting mechanisms

The effect of variations in the DAP current are particularly hard to predict because several other currents overlap with the depolarization from dendritic Nav channels. K channels, muscarinic or 5HT receptors, and GABAergic inputs can influence the after-potential (Marquez et al., 2013). In particular, somatic spikes are followed by both fast and slow AHPs that help repolarize the cell and lengthen the interval to the next spikes (Turner et al., 2002). Therefore, the DAP and the AHP have opposite influences on bursting. ELL expresses two different subtypes of SK channels (SK1: dendrites; SK2: soma) that cause AHPs (Ellis et al., 2007b, 2008) following a gradient where LS has a denser distribution than CMS. The relatively short DAP (8-10 ms) temporally overlaps with the longer AHP and their strength varies with PC subtype. It is possible that higher expression of dendritic Nav channels may partially compensate higher hyperpolarizing currents and allow these currents to be modulated with different gains.

Several mechanisms for external modulation of these hyperpolarizing and depolarizing currents have been characterized. Inhibitory interneurons are prevalent in the ELL (e.g. VML cells or granular cells) and synapse both on PC’s soma and dendritic arbor (Berman and Maler, 1999). Dendritic application of GABA_A_ agonist can affect the dendritic leak conductance leading to a DAP reduction and thus have a divisive effect on the cell’s input-output relationship (Mehaffey et al., 2005). In contrast, somatic inhibition has a subtractive effect in suprathreshold regime and divisive in subthreshold regime since the reversal potential of GABA_A_ channels is close to the neuron’s resting potential (Doiron et al., 2002). Therefore, somatic inhibition could also potentially interact with the subthreshold dynamic influenced by the DAP and AHPs. Since the inhibitory surround input onto pyramidal cells varies across segments (Hofmann and Chacron, 2017) and the proportion of inhibitory interneurons (supplemental Figure 3) also varies across segments, GABAergic inhibition is another segment-specific factor that could interact with the current from dendritic Nav channels.

5-HT is an important modulator of social behavior that is released during communication and LS shows the highest 5-HT innervation (Johnston et al., 1990). Interestingly, the innervation pattern is also layer specific across segments, where LS shows dense expression in the pyramidal cell layer and VML. 5-HT increases burst firing across segments with the greatest effect in the LS which is consistent with its expression density (Deemyad et al., 2011). These effects are mediated by 5-HT2 receptors (Larson et al., 2014) that increase PC excitability and bursting via downregulation of SK channel and M-type potassium channel currents that contribute to the AHP (Deemyad et al., 2013).

Given that several factors affect the shape of spike’s after-potential and the bursting dynamic we cannot be certain that the shorter burst ISIs we observed in LS are due to the denser Nav expression but it is a plausible factor. A strong DAP, that peaks a few ms after the spike, could explain the relatively high probability of having 2-4 ms ISIs in LS whereas the longer-lasting AHPs strongly influencing LS ON-cells could lead to the relatively low probability of ISIs longer than 5ms. The longer burst ISIs observed in CMS could result from the lower Nav current since it can delay the DAP (Fernandez et al., 2005) and is not opposed by a strong AHP in superficial PCs. We therefore propose that differences in Nav channel expression interact with other aspects of the neuron’s dynamic to influence the spiking patterns.

### Bursts and neural coding

Bursts have a well-defined role in coding for specific temporal features of sensory signals. The relationship between patterns of spikes within a burst can further signal details about the feature that triggered the burst (Oswald et al., 2007). Specifically, the ISI within the burst is correlated with the amplitude and slope of the upstroke it signals. Studies in several sensory systems have shown that burst structure can carry information about the stimulus (Guido et al., 1995; Lesica and Stanley, 2004) and that this information is behaviorally relevant (Marsat and Pollack, 2010). Our results point to the possibility that variations in Nav channels may influence the burst dynamic to adjust the correlation between burst ISI and stimulus features. By adjusting the range of ISI coding for specific portions of stimulus space, each ELL segment could have its burst-code adjusted for slightly different stimulus features. We already know that bursts are involved in processing different signals across segments since bursts signal the presence of certain types of chirps in LS but not the other segments (Marsat et al., 2009). Also, bursts in CLS and CMS might be more specifically dedicated to signaling prey-like stimuli (Nelson and Maciver, 1999). Differences in Nav expression we describe could contribute to these different roles of burst coding across segments.

We started this article by pointing out two perspectives on neural heterogeneity. One highlights how similar neural outputs can arise from diverse combinations of neural properties. The other focuses on the need to vary neural physiology in order to adjust and specialize cells for different purposes. Our study does not determine what role the variability in dendritic Nav channel expression plays: compensating for other neural properties to keep burst coding functioning similarly across segments or on the contrary adjusting burst coding for specific roles. Since these two possibilities are not exclusive, we speculate both may be at work. The differences we describe in our study could partly compensate for other variations in PC properties while at the same time influence burst structure and coding to improve the ability of different segments to perform specific tasks.

## Acknowledgements

We thank Dr. Andrew Dacks and Dr. Saravanan Kolandaivelu for offering their time, resources and guidance to help with some of the experimental procedures. Dr Michael Markham and Dr Vielka Salazar and their lab members were helpful in establishing the immunolabelling protocol. We also thank Dr Maurice Chacron and Dr Len Maler for useful discussions about this project. This research was supported by an NSF (IOS # 1557846) and West Virginia University Research Corporation.

## Contributions

GM designed the research, supervised the experiments and data analysis and wrote the paper. SIM performed the Nav immunolabeling experiments and analyzed them and wrote the paper. KMA performed the GABA immunolabelling experiments and analyzed them and wrote the paper. DKW performed the electrophysiological recordings and analyzed them.

## Supplemental Material

**Supplemental figure 1.**
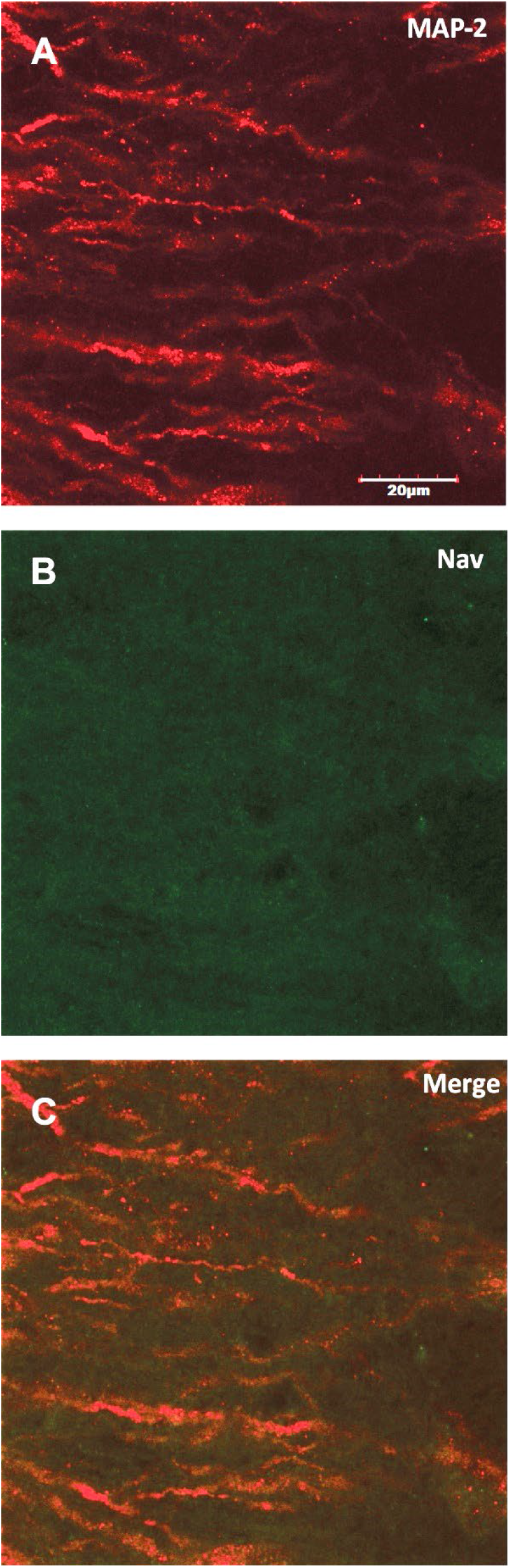
Absorption control: Confocal scan showing apical dendrites stained by Anti-MAP2 marker **(A)** in the ELL-LS segment. Anti-pan Nav antibody was incubated with immunogen (peptide) in a 1:10 ratio overnight on a nutator at 4 degrees. Slide mounted ELL sections were incubated with pre-adsorbed Nav antibody along with Anti-MAP 2 primary antibody which was followed by incubation with their individual secondary antibodies. No Nav expression was observed **(B,C)** demonstrating the specificity of Nav antibody used. Scalebar −20 μm (60X objective, **2X** digital Zoom). All scanning parameters and other steps of the protocol where identical to the ones used in the results section. Note that brightness of the image was increased to see the autofluorescence otherwise the image would have been completely black.

**Supplemental Figure 2.**
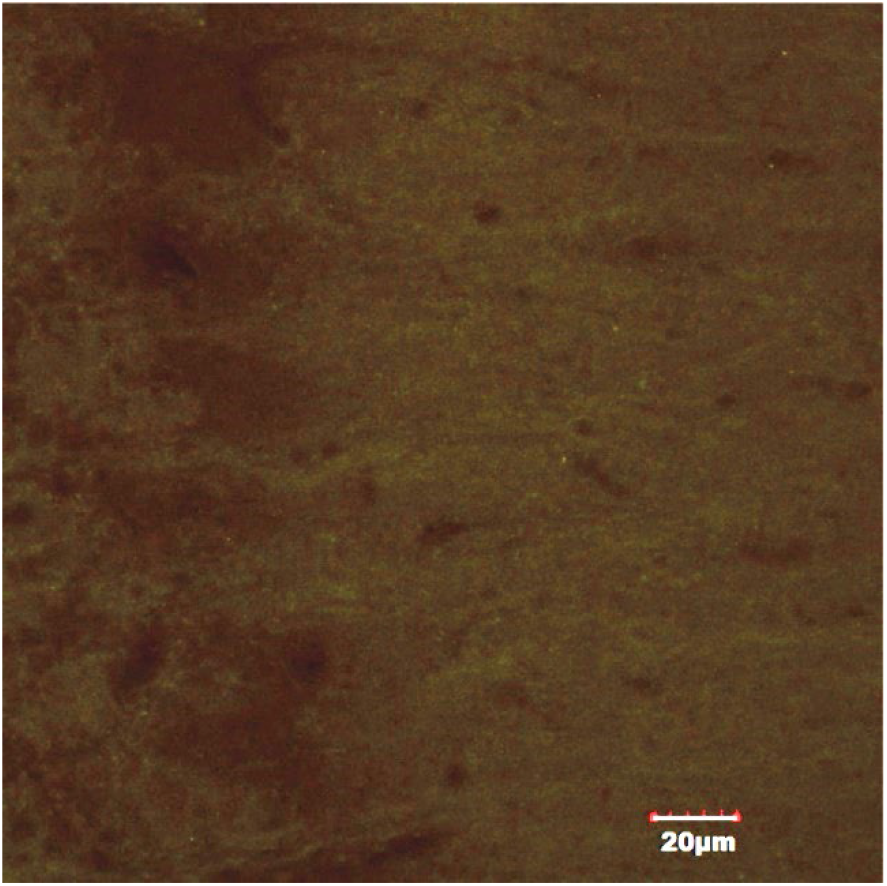
No primary antibody control: Anti-pan Nav and Anti MAP-II antibody used for labelling Nav and microtubules respectively were omitted from the IHC protocol as a control for non-specific binding of the secondary antibodies. Scalebar −20 μm (60X objective, **1X** digital Zoom). Note that brightness of the image was increased to see the autofluorescence otherwise the image would have been completely black.

**Supplemental Figure 2:**
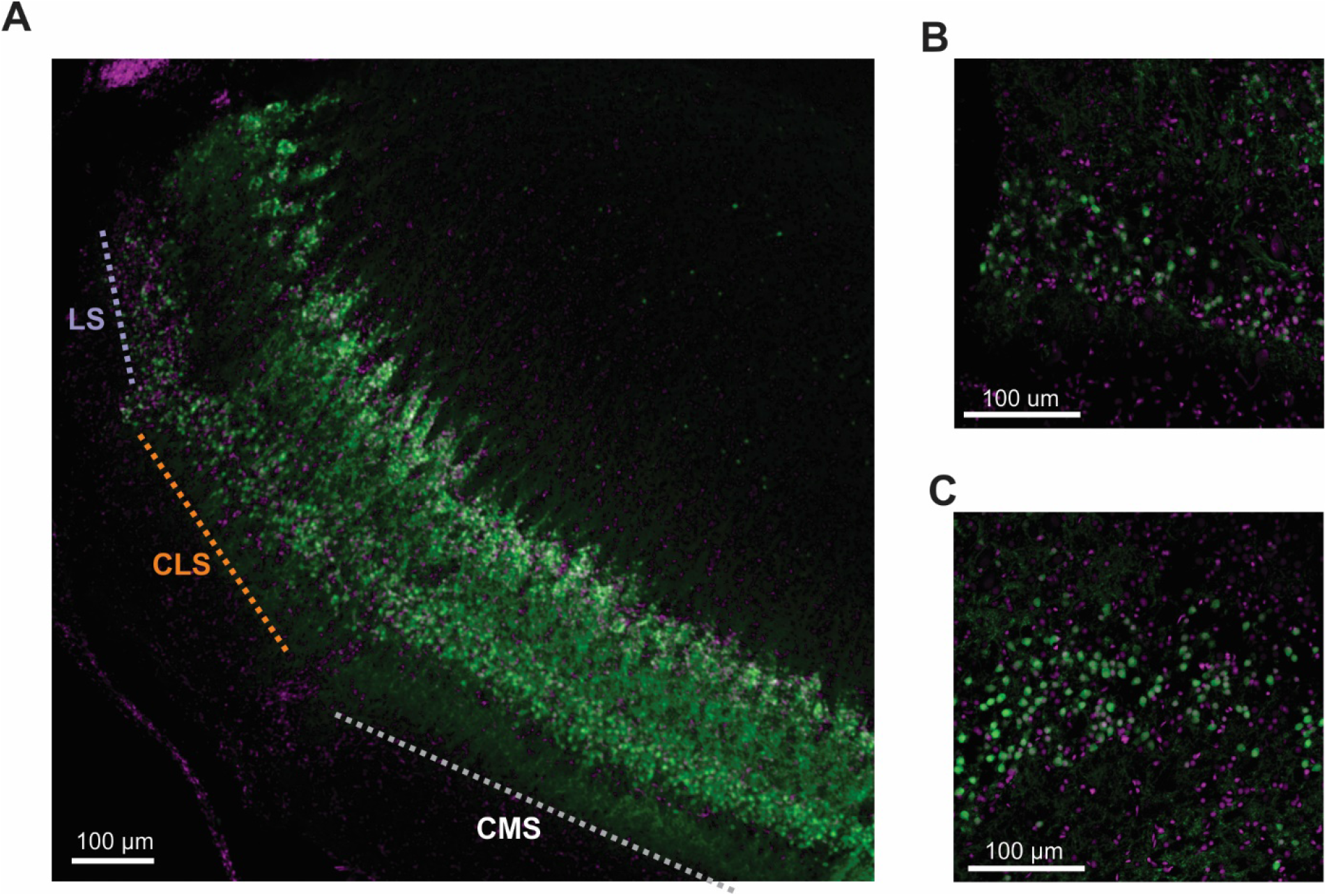
Gradient across ELL segments in number of GABA-labeled cell bodies. **A.** As described in the discussion, GABA may influence spiking dynamics. Preliminary data shown here (N=1 animal, n=4 sections sampled) indicate that the distribution of GABAergic interneurons vary across ELL segments. 10x magnification of ELL stained with Sigma Aldrich Anti-GABA produced in rabbit antibody (Sigma Aldrich # A2052) with Goat anti rabbit Alexa 488 secondary (Life Technologies, #A-11008; green) and a nuclear marker (Syto 59, Lifetechnologies #s11341; magenta) show a gradient of GABA distribution across segments. **B.** 40x magnification of LS and CLS (C). Only LS and CLS data have been quantified. We counted the number of GABA-labeled cell bodies in the granular cell layers in 100μm x 100μm area. Overall differences in mean number of interneurons were not found (LS=161.77 CLS=193.83, p=.52) but the variation in interneurons found was much higher for CLS, suggesting that the upper limit of interneurons found in this section is higher (standard deviations: LS=47.2726, CLS=137.5361)

